# Pathogen-microbiome interactions and the virulence of an entomopathogenic fungus

**DOI:** 10.1101/2023.12.18.572164

**Authors:** Matthew R. Kolp, Yazmin de Anda Acosta, William Brewer, Holly L. Nichols, Elliott B. Goldstein, Keertana Tallapragada, Benjamin J. Parker

## Abstract

Bacteria shape interactions between hosts and fungal pathogens. In some cases, bacteria associated with fungi are essential for pathogen virulence. In other systems, host associated microbiomes confer resistance against fungal pathogens. We studied an aphid-specific entomopathogenic fungus called *Pandora neoaphidis* in the context of both host and pathogen microbiomes. Aphids host several species of heritable bacteria, some of which confer resistance against *Pandora*. We first found that spores that emerged from aphids that harbored protective bacteria were less virulent against subsequent hosts and did not grow on plate media. We then used 16S amplicon sequencing to study the bacterial microbiome of fungal mycelia and spores during plate culturing and host infection. We found that the bacterial community is remarkably stable in culture despite dramatic changes in pathogen virulence. Last, we used an experimentally transformed symbiont of aphids to show that *Pandora* can acquire hostassociated bacteria during infection. Our results uncover new roles for bacteria in the dynamics of aphidpathogen interactions and illustrate the importance of the broader microbiological context in studies of fungal pathogenesis.

## Introduction

Fungal pathogens interact with hosts within complex communities of other microbes, and bacteria have been shown to influence host-pathogen dynamics in several ways. Host-associated bacteria can confer protection against fungi (e.g. in insects (1, 2), crustaceans (3), plants (4), reptiles (5), and humans (6, 7)). In turn, there is increasing evidence that fungal pathogens are also hosts to bacterial communities that can affect the severity of disease (i.e. virulence). For example, the fungal pathogen *Rhizopus oryzae*, which causes rice seedling blast, relies on toxin-producing bacteria that live within the fungus to successfully infect hosts (8). In contrast, a study of *Fusarium oxysporum* showed that a community of surface bacteria limited the ability of a fungal isolate to infect its plant host, and removal of the bacteria restored virulence (9). Recent studies suggest that the capacity to associate with bacteria is likely widespread among fungi (10-12), and therefore, studying how bacteria shape host-fungal pathogen interactions is critical (13).

The order Entomophthorales (phylum Entomophthoromycota, Humber) includes hundreds of species of fungi that are pathogenic to insects (14-16) (and in some isolated cases, humans (17)). These fungi play important roles in the population dynamics of insect hosts and could potentially serve as effective biocontrol agents (18, 19). Infectious fungal spores invade new hosts, and fungi then multiply as hyphal bodies within an insect. Hyphae then produce structures called conidiophores that penetrate and elongate through the cuticle of a host surface and then forcefully release asexual spores called conidia (20). Much about the biology of the Entomophthorales is unknown, including whether these fungi typically associate with bacteria.

*Pandora neoaphidis* (Remaudière and Hennebert; hereafter ‘*Pandora’*) is an important species of Entomophthorales that infects aphids (Hemiptera: Aphidoidea) and plays a role in the population dynamics of these crop pests (21-24). Previous studies have found that natural isolates of *Pandora* infect aphids at different rates (25, 26), but little is known about the factors shaping the virulence of this pathogen. However, a recent study used RFLP markers and microscopy to show that there are bacteria associated with *Pandora* and that microbial communities varied across isolates with different levels of virulence against hosts (27). From working with this pathogen in the lab, we have also observed that *Pandora* isolates rapidly lose virulence after subculturing on solid media. This is a phenomenon referred to as ‘degeneration,’ which is typical of species in the Entomophthorales (28). We hypothesized that changes in fungal microbiomes could be associated with degeneration in the lab.

After infecting a host, *Pandora* encounters a community of aphid-associated bacteria. These microbes include several species of ‘facultative’ bacteria that are not required for host survival but have different phenotypic effects on hosts (29). Several distantly related species of bacterial symbionts, including *Regiella insecticola*, confer protection to aphids against *Pandora* (30-32) (but not against a generalist species in the order Entomophthorales (33)). However, the protection is not perfect and pathogens are able to overcome symbiont-mediated protection in both lab infections and in the field (34). It is unknown if protective symbionts have any effect on fungal pathogens after infection. But, because aphids reproduce asexually in the summer and are typically surrounded by genetically identical offspring, any effects of aphid symbionts on fungal virulence against subsequent hosts could have important effects on disease dynamics in this system.

We studied the virulence of *Pandora* in the context of the wider bacterial community associated with both the aphid and fungal pathogen. We first measured whether fungal spores emerging from an aphid differ in virulence depending on the host’s microbiome. Spores emerging from aphids harboring a protective facultative symbiont were less virulent against subsequent hosts and did not grow when cultured on plate media. We then used 16S amplicon sequencing to characterize the bacterial microbiome of *Pandora*. We cultured fungal spores on plate media, quantified changes in virulence over subsequent plating, and found that the bacterial microbiome associated with *Pandora* mycelium is stable across multiple plate passages despite the loss of virulence. When we re-infected aphids with spores produced in culture, we found that *Pandora* regained virulence, but the bacterial community was still largely unchanged. Finally, we used an experimental manipulation to show that fungal spores can acquire bacteria from aphids during infection. Together our results highlight the importance of the wider bacterial community context for animal host–fungal pathogen interactions.

## Results

We tested the effects of the protective bacterial symbiont *Regiella insecticola* on the virulence and growth of the fungal pathogen *Pandora neoaphidis*. We used a panel of genetically identical aphid lines that each harbored a facultative symbiont or no facultative symbiont (control). This included two strains of *Regiella insecticola*, one that confers strong protection against *Pandora* (strain .313) and one which confers little to no protection (strain .515) (35). We also included aphids harboring *Serratia symbiotica* (strain .509) which is not protective against fungal pathogens (29). We confirmed that symbiont background influences the percent of exposed aphids that produced a sporulating cadaver (F(3) = 23.8, p < 0.0001; Figure 1A). This effect was driven by the ‘protective’ strain of *Regiella* (strain .313) that decreased the percent of aphids that died and produced sporulating cadavers compared to symbiontfree controls (strain .313, z = -6.6, p < 0.001). The other two symbionts had no effect on fungal protection.

**Figure 1:**
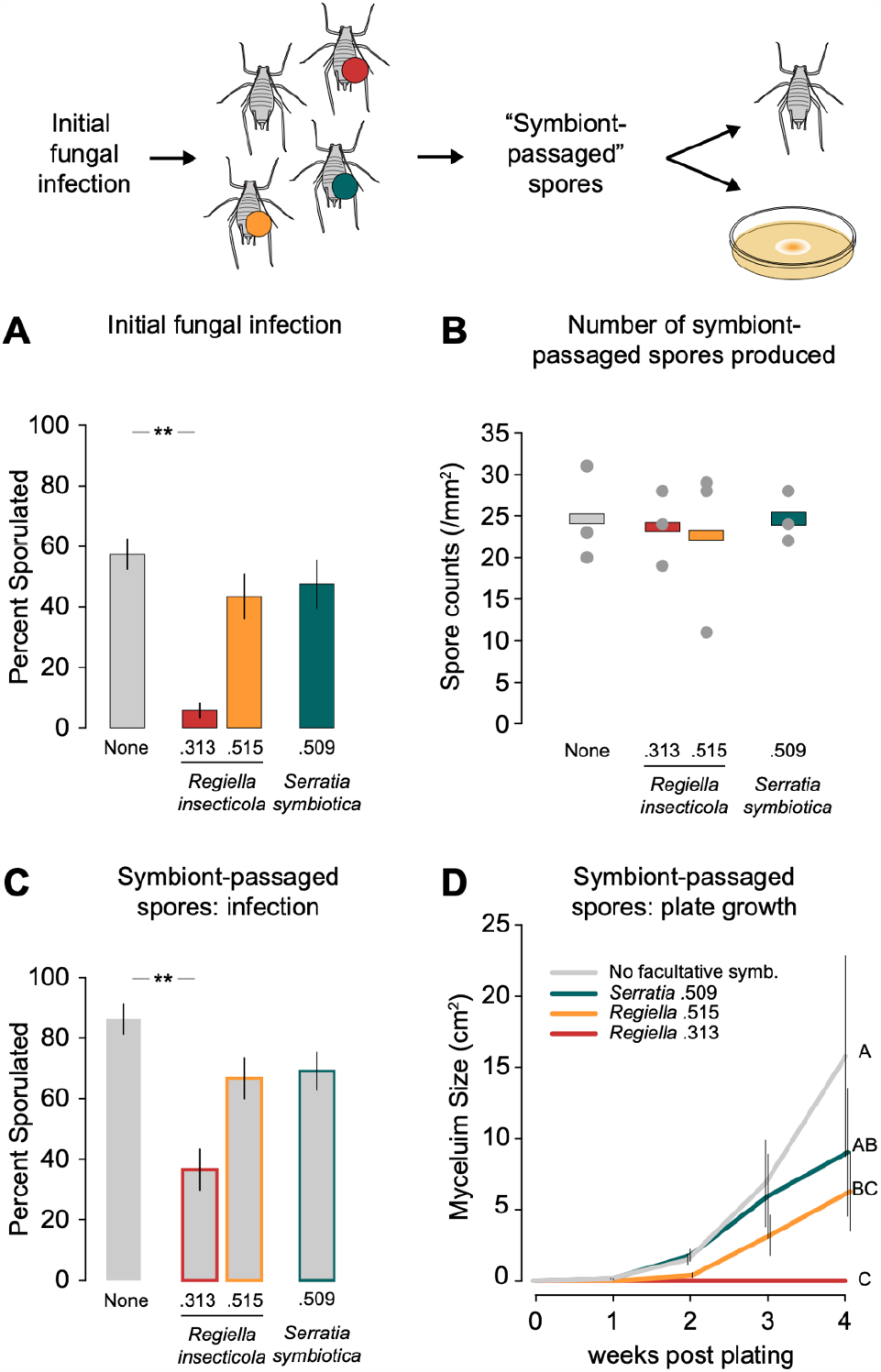
Effect of aphid facultative symbiont on fungal virulence and plate growth. An illustration of the experimental design is shown along the top of the figure. **A:** Percent of aphids that became infected with *Pandora* and produced a sporulating cadaver in the initial fungal infection. Facultative symbionts in each aphid line are shown along the bottom of the figure. The error bars show +/-one standard error, and statistical significance (**; p < 0.01) is shown along the top. This initial fungal infection was used to produce ‘symbiont passaged spores’ that emerged from infected aphids (i.e., cadavers) with different facultative symbiont backgrounds. **B:** Mean number of ‘symbiont passaged’ spores that emerged from cadavers (gray dots) produced in the initial infection as quantified under a light microscope per mm^2^. **C:** The percent of symbiont-free aphids that became infected when exposed to symbiont-passaged spores from each treatment. **D:** Mean size of mycelium growth from symbiont-passaged spores plated on media. The y-axis shows mycelium size (cm^2^), and the x-axis shows the number of weeks post initial plating. Statistically significant groups determined by post-hoc analysis were identified at four weeks post plating and are shown to the right of the figure. Error bars show +/-one standard error.

Next, we collected the aphid cadavers resulting from fungal infections of each of these lines, induced cadavers from each line to produce *Pandora* spores, and counted the spores produced by each cadaver. Symbiont background had no effect on the number of spores produced by cadavers (ANOVA, F(3,8) = 0.067, p = 0.98; Figure 1B). To test the virulence of these ‘symbiont-passaged’ spores, we used them to infect symbiont-free aphids. We found that the cadaver’s symbiont background explained the virulence of symbiont-passaged spores towards symbiont-free aphids (F(3) = 9.70, p < 0.0001; Figure 1C). Specifically, spores passaged though aphids harboring ‘protective’ *Regiella* strain .313 were less virulent against subsequent hosts spores passaged through symbiont-free aphids (strain .313, z = -4.8, p < 0.001). We found no effect of the other two symbiont strains (strain .515, z = -2.3, p = 0.058; strain .509, z = -2.1, p = 0.094). We also plated symbiont-passaged spores on media and found that symbiont background influenced mycelium growth on plates (Generalized Linear Mixed Model, symbiont χ^2^ = 31.7, df = 3, p < 0.0001). In this assay, passing through a host harboring either strain of *Regiella* significantly decreased fungal growth when compared to spores produced by symbiont-free aphids (strain .313: z = 5.8, p < 0.001; strain 515: z = 3.4, p = 0.0036). After 4 weeks of observation, there was no growth from spores exposed to aphids with *Regiella* strain .313 (Figure 1D).

Anecdotally, we have noticed that *Pandora* isolates rapidly lose virulence toward aphids after culturing the fungus on solid media. Our next objective was to determine if this loss in virulence is associated with changes in *Pandora’s* bacterial microbiome, and to determine whether host infection changes the associated bacterial community. An important methodological detail of this study is that we are not able to precisely control the dose of spores that are produced during an infection. Instead, we performed a ‘low-dose’ assay by exposing experimental aphids to sporulating fungus for one hour, or a ‘high-dose’ infection for five hours, and we then quantified the number of spores used in the infections. The *Pandora* spores used to inoculate the initial plate culture were highly virulent against symbiont-free aphids at a low-dose infection (1-hour infection; 18 spores/mm^2^; 96.7% of aphids infected; Figure 2A). We used some of these spores to establish a panel of plated cultures of *Pandora*, and we grew each isolate on solid media. After 1 month, we transferred small pieces of mycelium to fresh plates.

**Figure 2:**
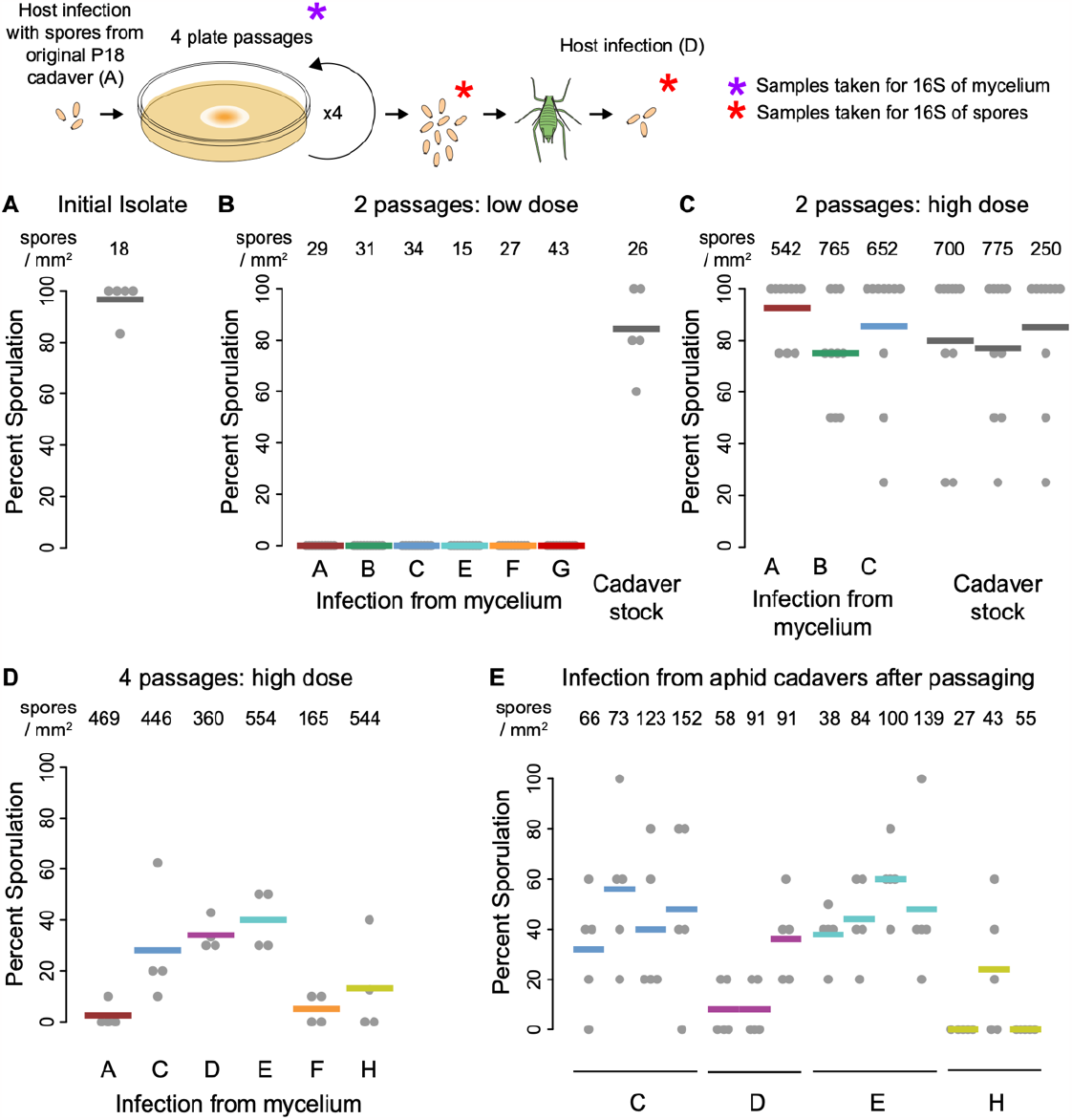
Aphid infections during the plate passaging experiment. An illustration of the experimental design and sample collection is shown along the top of the figure. **A:** Percent of symbiont-free aphids that became infected and produced a sporulating cadaver at a low spore dose infection using spores produced from an aphid cadaver. Spores were also used to inoculate initial plates for the plate passaging experiment. **B:** Percent of aphids that became infected at a low spore dose after 2 plate passages. The spore doses for each plate passaging replicate line are shown along the top of the figure. Plate replicate line (letters) is shown along the bottom of the figure, with the average of each shown with a different colored bar. An infection with *Pandora* maintained through re-infecting aphids (i.e., cadaver stock aphids) instead of culturing on plates is shown to the right of the figure. **C&D:** Aphid infections using high-dose assay of *Pandora* spores at 2 plate passages and 4 plate passages, respectively. **E:** Aphid infection with spores from aphid cadavers produced from plate passaging *Pandora* using the low-dose assay.

After two of these plate passages, spores produced in culture failed to infect any aphids at the lowdose infection (1-hour infection; 15 – 43 spores/mm^2^; 0% infected; Figure 2B). Spores produced in parallel by cadavers from aphid passaging were still virulent against hosts (1-hour infection; 26 spores/mm^2^; 85% infected; Figure 2B). We then repeated the infection with a high-dose (5-hour infection; 542 – 764 spores/mm^2^) and found that the high-dose of spores produced from culture plates was able to infect hosts (88% - 100% infected; Figure 2C). However, after four plate passages, virulence of spores produced from culture plates was further reduced with the high-dose assay (5-hour infection; 165 – 554 spores/mm^2^; 3% - 39% infected; Figure 2D). Of the few cadavers produced from this infection experiment, we found that the spores that emerged from this infection had, to some extent, regained virulence. After passaging through an aphid, spores were again able to infect aphids at the low spore dose (1 hour infection 27 – 152 spores/mm^2^; 0% - 60% infected; Figure 2E).

During the plate passaging and infection experiments (Figure 2), we collected samples for 16S amplicon sequencing. We included mycelium cuttings from replicate plate lines at each passage: the initial cultures through the fourth plate passage. We also sampled spores produced from the fourth plate passage just before aphid infection (Figure 2D) and the spores produced from re-infected aphids that did not require a high-dose of spores to successfully infect symbiont-free aphids (Figure 2E). Sequencing yielded 12,536 OTUs after removing non-bacterial reads. We removed OTUs from our dataset that were not present in at least three samples with more than ten sequence reads, leaving 369 OTUs after thresholding (86% of the 1,395,421 total reads were retained). We used these data to address two questions about the microbial communities associated with *Pandora*.

First, we analyzed the community of bacteria associated with fungal mycelium at each of the four plate passages. Bacterial richness did not change during plating (F_4,31_ = 2.51, p = 0.061). Further, we found no difference in alpha diversity across plates as measured by Shannon (p = 0.2798) and Simpson (p = 0.4447) diversity measures. Community composition did shift during subculturing *Pandora* (Figure 3A; F_2,24_ = 1.54, p < 0 .002), and the stage of plate passage explained 16% of the variation in bacterial community composition (R^2^ = 16.1%). However, we found no evidence that any individual OTUs differed between any of the plate passages based on differential abundance testing. Our analysis also included a block effect for the line of plate subculture (e.g., line A-H for plate passage 1-4), which explained more (23%) of the community wide variance. Taxonomically, mycelium samples of *Pandora* were associated with four main phyla of bacteria, which made up 97% of 16S reads (Figure 3B): Proteobacteria (55.6% of reads), Firmicutes (20.9%), Actinobacteriota (15.1%), and Bacteroidota (4.7%) (Figure 3B).

**Figure 3:**
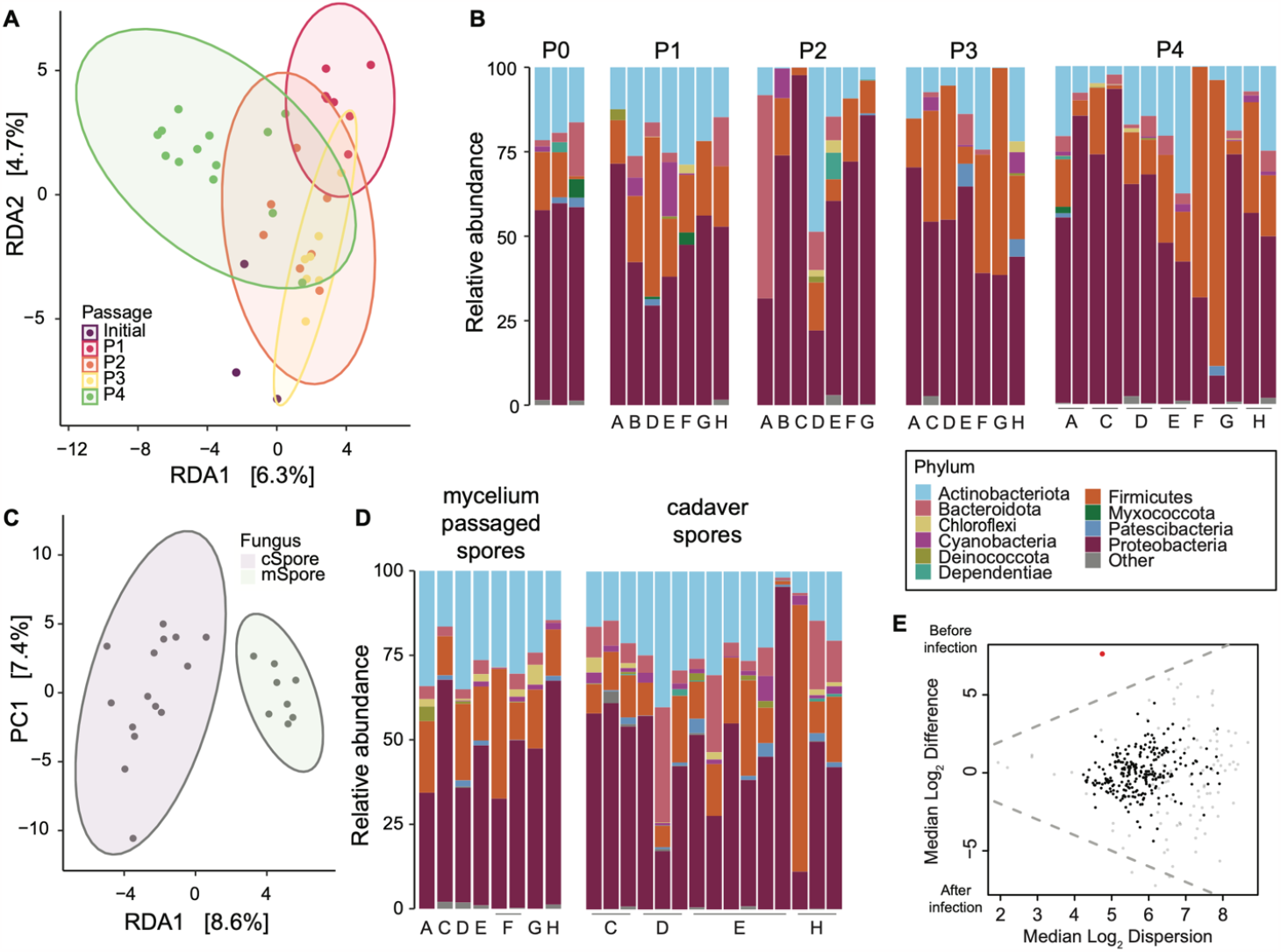
16S analysis of *Pandora*-associated bacterial communities. **A**. Adonis permutation and constrained ordinations (CAP Euclidean) plot by plate passaging. Redundancy analysis for bacterial communities from monthly plate passage (P# by color) of *Pandora* through laboratory culturing on Petri plates containing ‘ESMAY’ media. **B**. Phylum-level bar plots of taxonomy change by plate passaging. Taxonomic barplots for bacteria associated with mycelium across plate passage (P#) of *Pandora* through laboratory culturing on Petri plates containing ‘ESMAY’ media. Each sample from a replicate line is represented by a single bar with the relative abundance (non-CLR transformed data) of the top 10 phyla. **C**. Adonis permutation and constrained ordinations (CAP Euclidean) plot by spore type. Redundancy analysis for bacterial communities associated with spores collected from the fourth plate passage (mSpores) and those generated after reinfecting aphids (cSpores) with mSpores. **D**. Phylum-level bar plots of taxonomy change by spore type. Taxonomic barplots for bacteria associated with spores collected from the fourth plate passage (mSpores) and those generated after reinfecting aphids (cSpores) with mSpores. Each sample from a replicate line is represented by a single bar with the relative abundance (non-CLR transformed data) of the top 10 phyla. **E**. Differential abundance plot. Effect plot showing the relationship between Difference and Dispersion for all OTUs included in analysis (n=369). The red dot (OTU 27; Proteobacteria, Neisseraceae) represents a differentially abundant OTU (Welch’s test); grey dots are abundant, but not differentially abundant; black are rare, but not significantly rare.

We then compared the bacterial community associated with *Pandora* spores before (plate-passaged ‘mycelium’ spores) and after aphid re-infection (‘cadaver’ spores). To do this, we induced mycelium to produce spores after the four plate passages by placing mycelium cuttings on TWA. We used these spores to infect aphids and compared them to the spores that emerged from successfully infected aphids. We found that the richness (F_1,21_ = 0.06, p = 0.81) and diversity (Shannon – F_1,21_ = 1.18, p = 0.29; Simpson – F_1,21_ = 0.98, p = 0.33) did not change after re-infection of the host (Figure 3C). Community composition of bacteria associated with spores did change with aphid re-infection (Figure 3D; F_1,21_ = 1.89, p < 0 .001), with spore sample type explaining 8.4% of the variation in community composition. Furthermore, only one OTU (OTU 27; Proteobacteria, Neisseraceae [uncultured]) changed in abundance with aphid infection after implementing a Benjamini-Hochberg correction. This taxon became less abundant with host infection (Figure 3E). Of the main four bacterial phyla associated with *Pandora* mycelium, Bacteroidota increased from an average relative abundance of 2.8% in spores generated after plate passaging to an average of 9.4% in spores collected after re-infecting aphids. The other three main phyla changed only slightly in their relative abundances (Proteobacteria (mSpore_mean_ = 57% vs. cSpore_mean_ = 47%), Actinobacteriota (26% vs. 21%), and Firmicutes (19% vs. 17%).

We next used an experimental approach to test whether pathogens can acquire specific host-associated bacteria from aphids during infection. We chose a microbe found in the aphid gut that can be genetically manipulated in the lab and grown *in vitro* (36-38). We were able to transform *Serratia symbiotica* strain CWBI 2.3T using mini-Tn7 site-directed transposon insertion. Transformed bacteria were able to grow on plates of Trypticase Soy Broth/Agar (TSB) with zeocin (InvivoGen). In this experiment, we infected aphids with *Pandora* (from cadaver stock aphids) and then injected them with transformed *Serratia*. Control aphids were injected with transformed *Serratia* but were not exposed to *Pandora*. All aphids died between 4-6 days after injection. Dead aphids from the control and cadavers produced from *Pandora* infection were then placed over liquid media to sporulate, which was then spread plated on replicate plates of TSB + zeocin for 48 hours at 27°C. No plates from control aphids grew bacterial colonies; however, *Pandora* spores from infected aphids successfully transferred the transformed *Serratia* to plates at varying CFUs (Figure 4).

**Figure 4:**
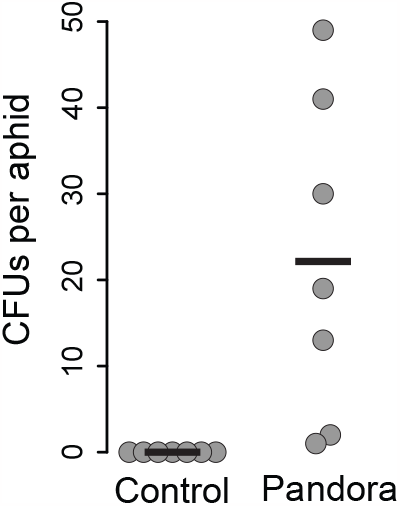
Fungal acquisition of host-associated bacteria. An illustration of the experimental design is shown starting on the left. The figure on the far right shows the number of CFUs resulting from control and *Pandora-*infected aphids spread-plated on replicate plates of TSB + zeocin after 48 hours. Each replicate is shown with a grey point, and means are shown with a black bar.

## Discussion

Aphid facultative symbionts confer protection against specialist fungal pathogens in the family Entomophthorales (30-33). We found that when a fungal pathogen is able to overcome this protection and infect a host (which occurs in both the lab and field (34, 35, 39)), the pathogen suffers reduced virulence against subsequent hosts. Specifically, we found that spores that emerged from aphids harboring a protective strain of *Regiella insecticola* were less virulent against subsequent hosts, and these protective *Regiella-*passaged spores did not grow on plates in the lab. Asexual, reproductive female aphids are typically surrounded by genetically identical offspring harboring the same heritable bacteria. The additional protective effects of *Regiella* that we uncovered likely benefit an aphid lineage even if the symbiont fails to protect an individual aphid. This finding has important implications for our understanding of aphid population dynamics, and for our understanding of the evolution of symbiontmediated protection against pathogens and parasites (40).

Pathogenic fungi commonly lose virulence and experience morphological changes when cultured on artificial media (termed ‘degeneration’ (28)). This phenomenon is a concern for the study of entomopathogens in the lab and for the use of fungal pathogens in biocontrol and other applications (reviewed in 28). We quantified degeneration of *Pandora neoaphidis* on media. We showed that plate culturing of *Pandora* in the lab is associated with a loss of virulence towards aphids, with pathogen spores unable to infect hosts at low doses after two plate passages. After four plate passages, *Pandora* virulence was further reduced at high dose infections. *Pandora* spores regained virulence after reinfection of the aphid host. These findings are relevant to the potential use of *Pandora* as a biocontrol agent (18, 19, 41, 42), and suggest that additional research into entomopathogen degeneration is needed before these biocontrol microbes could be grown at scale for use in agricultural pest management.

We hypothesized that changes in *Pandora’s* bacterial microbiome could explain degeneration (reduced virulence) on plate media. Contrary to our expectations, changes in fungal virulence do not appear to be associated with changes in the bacterial microbiome. Instead, we found that richness and diversity of the bacteria associated with *Pandora* changed very little during plate passaging and reinfection experiments. While the total community composition of these bacteria was influenced by the number of plate passages and through reintroduction of the pathogen to its host, most of the variance in these models was not explained by these factors. In fact, by including a block effect of replicate plate line in our plate passaging experiment, we found that the plate line explained a greater percentage of the variation in community composition in *Pandora* cultures. This also suggests that *Pandora* is getting a portion of its bacterial microbiome from its parental subculture, rather than the environment. Together our results suggest that a change in the bacterial microbiome of *Pandora* is not the specific epigenetic mechanism driving the loss of virulence observed in the lab. Future work should investigate alternative, loss-of-virulence mechanisms in entomopathogenic fungi, including the roles of mycoviruses (43), DNA methylation, or alterations in karyotype (chromosome polymorphisms) (28).

In certain fungal taxa, the importance of bacteria in pathogenesis is more apparent. For example, a bacteria (*Burkholderia rhizoxinica*) secretes a toxic virulence factor for the fungus (*Rhizopus microsporus*) against its plant host (8). When removed of this bacterium, the fungus is no longer pathogenic. Our preliminary characterization of bacteria associated with *Pandora* did not reveal any specific OTUs that explained loss or reacquisition of virulence towards aphids. Thus, it seems unlikely that any bacterial microbiome members of *Pandora* are impacting virulence towards its host. Instead, the microbiome of *Pandora* may be compromised of “hitch-hiking” bacteria (11) with expanded niches to increase survival after aphid infection. More research would be needed to determine what roles bacteria are playing in *Pandora’s* biology. We showed that the bacterial microbiome of *Pandora* is composed of four main phyla: Proteobacteria, Firmicutes, Actinobacteriota, and Bacteroidota. A key limitation of our study is that we used only one *Pandora* genotype to test the relationship between virulence and the pathogen’s microbiome. We know that the pathogen genotype and the aphid symbiont genotype can interact to influence disease (35). Isolating additional fungal lines from natural populations could help identify ‘core’ bacterial taxa found across multiple fungal lineages.

Many entomopathogenic fungi emerge from their insect host’s body cavity where they encounter hemolymph, gut, and cuticle microbial associates. We last showed that *Pandora* acquired experimentally-injected gut bacteria in aphids that were viable after pathogen infection and dispersal from the host. We used a genetically modified gut symbiont of aphids engineered to encode for antibiotic resistance, and we demonstrated that we could culture this bacterium by plating fungal spores that emerged from an infected aphid. Our expectation is that transformed *Serratia* were transferred to media on the surface of *Pandora* spores. A previous study of the bacteria associated with *Pandora* suggested that some microbes can also be found inside *Pandora* cells (demonstrated through Fluorescence in situ hybridization and confocal microscopy (44)). Whether bacteria are encapsulated by *Pandora* during infection, or bacteria are attached superficially as their host becomes infected and dies, or both, is unknown. Current research suggests that fungi can associate with bacterial symbionts of animal and plant hosts in non-specific ways, and that these interactions are products of stress and environmental conditions (45). One interesting possibility is that bacteria associated with aphids could be transferred via *Pandora* to new aphid hosts.

Our study demonstrates two new ways that host microbiomes influence fungal pathogens: the microbiomes of infected hosts can influence virulence against subsequent infections, and hosts and fungi can potentially share bacterial taxa. Although we found no apparent link between the pathogen microbiome and virulence or growth in culture, we identified a relatively stable consortium of bacteria that associate with *Pandora*. Our results yield important considerations for future studies of *Pandora neoaphidis* and other fungal entomopathogens, how microbes shape pathogen evolution, and for the potential use of fungi in insect biocontrol.

## Methods

### Fungal isolate

We isolated a strain of a fungal pathogen that we visually identified as *Pandora neoaphidis* (referred to here as P18) from an unwinged adult pea aphid feeding on *Vicia sp*. in May of 2019 in Knoxville, TN, USA. We brought live aphids into the lab and harbored them on lab-grown *Vicia faba* (var: Windsor) plants at 20°C. We maintained aphids at ambient humidity (lab conditions), which causes entomopathogen-infected aphids to produce a dry ‘resting cadaver’ that does not release spores. We then selected a single resting cadaver and placed it on 2% tap water agar (TWA) overnight, which induces the cadaver to release spores (20). We established the fungus in the lab using two methods: culturing on plated media and aphid passaging (see below).

### *Pandora* plate culturing

We collected P18 spores using a sterile pipette tip from around the infected cadaver and inoculated the center of a Petri dish 10 cm^2^ containing ‘ESMAY’ media (1% yeast extract, 1% peptone, 1.5% agar, 4% maltose, and 2 egg yolks per 100mL media). We grew plates wrapped in Parafilm in the dark at 20°C. *Pandora* mycelium grew at 20°C on a plate for one month, at which point fungal mycelium was transferred to new plates by cutting approximately 1 cm^2^ squares of mycelium, removing fungus from the media, and placing it on a fresh plate. We noted the parental line of each subculture through the four plate passages.

### Molecular species confirmation for strain P18

We confirmed the identification of P18 as *Pandora neoaphidis* using PCR amplification and sequencing. We cut a 1 cm^2^ piece of fungal mycelium from plate growth and extracted DNA using phenol-chloroform with an ethanol precipitation. We performed PCR using previously published primers and conditions (PnITS_F: GAATAGATTGTCTTTATAACTACGTGTAGA and PnITS_R: ACCAGAGTACCAGCATATCC); 30s at 98°C followed by 30 cycles at 98°C for 30s, 61°C for 20s, and 72°C for 2 min, with a final extension at 72°C for 7 min. (19). Following PCR, we sequenced the amplicon via Sanger sequencing in the forward and reverse direction. We used BLASTN to show that the resulting consensus sequence had 100% sequence similarity to published sequences for *Pandora neoaphidis* (e.g. HQ677587.1).

### *Pandora* passaging through aphids

We maintained *Pandora* isolate P18 in ‘aphid culture’ by serially passaging it through healthy aphid individuals and storing the dried resting cadavers at 4°C. We reared pea aphids from the LSR1-01 genotype (originally collected near Ithaca, NY in 1991 from *Medicago sativa*) (46). We confirmed the absence of any secondary symbionts using PCR (47), and subsequently maintained this aphid line in the lab on *Vicia faba* at 16L:8D at 20°C. From the initial P18 cadaver collected in the field, we inverted the sporulating cadaver over LSR1-01 adult aphids. We then moved exposed aphids to *V. faba* plants covered by unvented cages for 48 hours to keep aphids under relatively high humidity, and then moved aphids to new plants with a vented cup cage in an incubator with low relative humidity. Starting on the fourth day after exposure to P18, aphids began to produce new resting cadavers, which we collected until day 8 post-exposure. Resting cadavers were stored at 4°C along with packets of Silica gel for up to a month. We repeated this infection experiment each month to generate new cadavers for long-term maintenance of a P18 in the lab.

### Aphid symbiont effects on pathogen virulence

We used a panel of LSR1-01 aphids with different facultative symbionts to investigate if aphid symbionts affect the pathogen’s virulence in subsequent infections. The panel included a protective strain of *Regiella* (.313), a strain of *Regiella* from another aphid species (*Myzus persicae*) identified in previous work (35) that does not confer protection in pea aphids (strain .515, (48)), and a strain of *Serratia symbiotica* (strain .509) isolated from pea aphids from Knoxville in 2019. *S. symbiotica* is not known as a protective symbiont of pea aphids against fungal pathogens (28). For the experimental infection of LSR1-01 with these different symbionts and a nosymbiont control (Figure 1), we performed a fungal infection on the panel, and collected dried cadavers resulting from successful infections. We then grew new LSR1-01 (symbiont-free) aphids and exposed them to ‘symbiont-passaged’ spores from cadavers from the initial fungal infection.

To perform the fungal infections, we took cadavers that were stored at 4°C and placed them on 2% TWA plates overnight to induce sporulation. 10-day-old adult aphids were placed at the bottom of an infection chamber: a PVC tube (5cm x 3.2cm) painted on the inside with fluon (Insect-A-Slip, BioQuip Products, Inc. product #2871A) to prevent aphids from crawling up the chamber. A single sporulating cadaver was placed at the top of each chamber facing down to allow spores to shower onto the aphids (20). For each experiment, fungus is rotated among the infection chambers every few minutes to ensure an equal dose of spores across chambers. After infection, aphids were then placed on new *V. faba* plants. Each plant was assigned a random number to ensure data collection was blind to treatment. Vented cages were covered with parafilm “M” to increase humidity. After two days, aphids were moved to new *V. faba* plants in unvented cages at ambient humidity. We recorded aphid survival and signs of sporulation (i.e. cadavers) for eight days until aphids ceased to show new signs of sporulation.

We analyzed aphid infection data using Generalized Linear Models (GLMs) with a quasibinomial error and logit link function implemented in R v.4.2.2. The sporulation status of each aphid was modeled as a binomial outcome, and symbiont was included as a fixed effect. In the initial infection, we also included experimental replicate as a fixed effect. Minimal models were derived by removing the fixed effects and performing model comparisons using ANOVA and F-tests. Post-hoc analyses comparing levels within symbiont were performed using the ‘multcomp’ package. Spore production was analyzed using a one-way ANOVA with symbiont background as the explanatory factor after checking for normality using the *aov* function in R v.4.2.2. Plate growth was analyzed using a linear mixed effects model implemented using the ‘lme4’ package in R v.4.2.2. Symbiont background and timepoint were included as fixed effects, and plate replicate was included as a random effect in the model; the model structure accounted for the paired nature of the analysis by including the random effect of plate. Minimal models were derived by removing the effect of symbiont and then timepoint, and models were compared using ANOVA and chi-squared tests. Post-hoc comparisons analyzing the effect of symbiont were performed using the ‘multcomp’ package.

### Virulence of spores from mycelium and from cadavers

To assess the effects of repeatedly subculturing *Pandora* on virulence, we performed experimental infections of aphids using spores generated from cultured mycelium. For the assay, we cut ∼1 cm^2^ plugs of mycelia from the leading edge of each P18 culture and placed each plug on separate 2% TWA plates to induce sporulation.

We performed two versions of this assay, one that exposed aphids to a ‘low’ spore dose (1 hour of exposure), and one that exposed aphids to a ‘high’ spore dose (5 hours of exposure). This is because we are not able to precisely control the number of spores produced by a mycelium plug. We quantified each spore dose for each experimental group by counting spores by including an infection chamber containing only a glass cover slip in the rotation. We visualized each glass cover slip under a compound microscope after the infection and counted the number of *Pandora* spores in three random 1mm ^2^ fields of vision, averaging these values for each measurement. We carried out these infections using the initial plate isolate, after two rounds of plate passaging, and after four rounds of plate passaging. At two rounds of passaging, we carried out both a low and a high dose infection, and we also included spores produced by aphid cadavers that had been maintained in LSR-01 adult aphids (see above). At four rounds of passaging, we performed only the high dose assay.

After the fourth plate passage of P18, we re-introduced P18 to healthy adult aphids with spores from mycelium plugs (Figure 2). From this infection, we collected dried aphid cadavers and stored them at 4°C. We used some of these dried cadavers to infect aphids to measure virulence. We used other cadavers to generate spores for 16S sequencing.

### DNA extraction and 16S sequencing of *Pandora*

For mycelium samples, we cut a 1 cm^2^ piece of the leading edge of the fungal culture and stored it at -80°C. For spore samples, we placed mycelium or a resting cadaver on 2% TWA inverted over a microcentrifuge tube with lysis buffer. After 24 hours of sporulation, the tube was vortexed to suspend spores and stored at -20°C. We extracted total genomic DNA from *Pandora* using phenol-chloroform with an ethanol precipitation and stored the samples at - 20°C. We included a negative control extraction on molecular grade water. We quantified sample DNA using an ND1000 spectrophotometer (NanoDrop Technologies), standardized DNA to 7.5 ng/μL, and PCR-amplified from these samples using primer pairs 341F/785R that target the 16S V3 and V4 region (49). PCR reactions included Kapa high-fi master mix at 1X, primers at 200nM each, 20ng gDNA, and molecular grade water. PCR conditions were 95°C for 3 min, 34 cycles of 95°C for 30s, 58°C for 30s, and 72°C for 30s, followed by 72°C for 5 min. We purified the PCR product with Agencourt Ampure XP beads and indexed each sample with a unique combination of forward and reverse Nextera XT v2, set A indexes (Illumina) using Kapa HiFi master mix (Roche) in a reduced-cycle PCR. We again purified the indexed PCR product with Agencourt Ampure XP beads, and visualized and quantified the final libraries for quality control (using an Agilent Bioanalzyer). The libraries were sequenced at a final loading concentration of 4 pM using 275 paired-end reads on an Illumina MiSeq instrument with version 3 reagents and 10% PhiX spike-in on an Illumina MiSeq at the University of Tennessee Genomics Core.

### *Pandora* mycelium and spore 16S analysis

We analyzed bacterial communities associated with fungal mycelium and spores via 16S sequence data using Mothur (v. 1.38) (5) following the MiSeq standard operating protocol (6). First, we removed reads with ambiguous bases and mapped reads to the V3/V4 region of the SILVA reference database (positions 1044 through 43116 in database v. 138.1; www.arb-silva.de), accepting alignments with >99% similarity. Next, we removed chimeric reads using vsearch within mothur and reads that aligned to DNA from chloroplast, mitochondria, archaea, and eukaryotes. We removed sequences found in the negative control from all other samples and determined OTUs using “dist.seqs” and “cluster” commands. Taxonomy of each OTU was assigned using the ‘classify.otu’ command. Raw sequence data are available through the Sequence Read Archive with BioProject ID: PRJNA1054062. Relative abundance of taxonomic groups, richness (Chao1), and diversity indices (Shannon and Inverse Simpson) were calculated before the removal of OTUs not observed ≥10 times in ≥3 samples due to their diminishing utility in analyses and potential to be sequencing artifacts (50-52). 86% of reads remained after thresholding. Counts of OTUs were normalized via a centered-log-ratio (CLR) transformation using the ALDEx2 package (53, 54).

We fit a linear model for each alpha diversity metric to test the similarity of mycelium cultures and spores. We assessed community composition by permutational multivariate analysis of variance (PERMANOVA) and constrained partial redundancy analysis (RDA) with the condition of log(reads) added to remove the effect of variable read depth in each sample. For mycelium samples, we included the plate line as a block effect. For PERMANOVA, we used the adonis2 function from the vegan package (55) to model between-sample Aitchison distances (54). The RDA was fit and visualized using an ordination plot of the significant constrained axes (RDA) or unconstrained axes (Principal Components). The CLR-transformed abundance of each genus was modeled to test responses to *Pandora* plate passage number and spore type. Across both models and where appropriate, p-values were adjusted to correct for multiple comparisons using a Benjamini-Hochberg correction.

### Transposon insertion using mini-Tn7 into *Serratia symbiotica*

We obtained isolate CWBI 2.3T of *Serratia symbiotica* (NCBI tax id: 138074) from the DSMZ-German Collection of Microorganisms and Cell Cultures. This species was originally isolated from the black bean aphid *Aphis fabae* (37) and can be cultured on Trypticase Soy Broth (TSB) or Agar (TSA) at 27°C. We grew microbes harboring the mini-Tn7 plasmid and the pTns2 transposition components on LB (56, 57). We grew *S. symbiotica* in 5mL of TSB for two days and then spun cells down at 4000rpm for 10m at 4°C. We removed the supernatant and added 1mL of 300nM sucrose solution, vortexed the mixture, and then centrifuged again for 5 minutes. We repeated this wash step a second time, and then resuspended the cells into 300mL of 300nM sucrose. We standardized the mini-Tn7 plasmid and the pTns2 transposition components to 50ng/mL and then added 2mL of Tn7 and 6.9mL of pTns2 to the two treatment samples. The mix was vortexed and then electroporated. We then incubated the samples at 27°C with shaking for 16 hours and plated 10mL on TSA with zeocin for 3-4 days until colony growth was observed (36, 38, 56).

### *Serratia* gut infection and fungal infection

For *S. symbiotica* injections, we grew transformed bacteria in TSB with zeocin, and then spun the cells for 10m at 4000rpm and washed the cells with PBS, repeating this procedure three times. We then normalized to an OD of 1, and diluted 100-fold in pH 7 adjusted Buffer A (25 nM KCl, 10nM mgCl 2, 250 nM Sucrose & 35 nM Tris-HCl) (38). We used a Femtojet injector (Eppendorf) to inject recipient aphids with 1mL each, after which aphids were housed on plants in cages as described above.

## Supporting information

Raw_data

## Acknowledgements

Sequencing was carried out at the University of Tennessee of Knoxville Genome Sequencing Facility. This work was supported by US National Science Foundation (NSF) Grant IOS-2152954 to BJP, and NSF Grant DBI-2109440 to MRK. BJP is a Pew Scholar in the Biomedical Sciences, supported by The Pew Charitable Trusts.

## Data availability

Data has been included as a supplementary file. Raw sequence data from the MiSeq run have been uploaded to the NCBI Sequence Read Archive (SRA) with BioProject ID: PRJNA1054062 and BioSample IDs: SAMN38882051-SAMN38882110.

## Author contributions

Conceptualization: HLN & BJP; Formal analysis: MRK, WB, & BJP. Funding acquisition: MRK & BJP; Investigation: YdAA, WB, HLN, EGB, KT; Writing – original draft: BJP. Writing – review & editing: MRK, BJP. All authors approved the final version of the manuscript.

